# Epigenetic field defect discriminates normal tissues from healthy and tumor-bearing kidneys

**DOI:** 10.1101/2023.10.05.560844

**Authors:** Jurgen Serth, Christel Reese, Tatyana Hubscher, Bastian Hill, Michael Klintschar, Jorg Hennenlotter, Markus Antonius Kuczyk

## Abstract

**Background:** The detection of alterations in DNA methylation (DNAm) in normal human tissues may provide molecular insights for cancer development, the estimation of personal cancer risk, and a basis for early diagnosis of malignancies to reduce cancer lethality. Thus, whether DNAm allows discrimination between healthy normal and normal tumor-adjacent renal tissues is an important question.

**Methods:** DNAm of loci in the LINE1 repetitive sequence and *ANKRD34B, NHLH2, TBR1*, and *ZIC1* was measured in a total of 493 tissue samples representing 342 normal autopsy and 151 histopathological normal tumor-adjacent renal tissues by pyro-sequencing a total of 22 CpG sites.

**Results:** Unsupervised k-means clustering demonstrated a significant imbalance of tissue samples in three stable tissue clusters. Random forest classification demonstrated discrimination of normal and normal tumor-adjacent renal tissues with a median area under the ROC curve of 0.81 (p=2.7×10^−9^, diagnostic odds ratio 10.4). Variable importance analysis revealed CpG sites in LINE1 and *ANKRD34B* as the most important model contributors.

**Conclusions:** DNAm alterations in normal tissues allow detection of renal cancer with high diagnostic efficiency, defining a DNAm field effect in the kidney. The methylation signature may serve as a basis for an epigenetic DNAm clock permitting estimation of individual renal cancer risk.

## 1. Introduction

DNA methylation (DNAm) is a frequently observed and comparatively stable epigenetic alteration in human tissues that is associated with a number of malignant and non-malignant diseases. The functional link between DNAm and gene silencing in stem cells has led to the epigenetic stem cell model explaining the frequent detection of cancer-specific hypomethylation and hypermethylation in malignant cells [1]. Moreover, tissue- and disease-specific DNAm patterns could provide a diagnostic basis for cancer prediction, early detection, and prognosis, as well as estimating the therapeutic response [2]. Renal cell cancer (RCC) has a prevalence of 3% and 5% in women and men, respectively, and approximately 25% of patients are diagnosed with initial distant metastasis [3]. The most relevant risk factors for RCC identified in epidemiological studies thus far are age, obesity, and smoking [4,5].

It has been known for some time that DNAm in tissues increases with chronological age in association with the number of cell divisions [6]. However, the probability that the exposure of tissues to cancer risk factors also affects DNAm similarly increases with age, leading to the observation of cumulative cell-intrinsic and -extrinsic DNAm in tissues [2]. Consequently, only the comparison of normal healthy tissues (N) and normal tumor-adjacent tissues (adN) obtained from individuals exposed to risk factors allow differentiation of age-related and age-independent DNAm associated with risk factors. In practice, tissue sampling of N is largely limited to post-mortem autopsy samples, with the challenge of potential tissue and DNA degradation. However, DNAm analysis of postmortem mouse kidney tissues over a period of 4 days demonstrated no effect of DNA degradation on measured DNAm levels [7].

On the other hand, adN are likely exposed to extrinsic risk factors considering that the risk-event cancer has already occurred and, in principle, may even be subject to further alterations until detection and surgical removal of the corresponding primary tumors. Few analyses of DNAm have made use of normal renal tissues thus far. Using a qualitative approach, Waki et al. found, in normal postmortem renal tissues, age-dependent DNAm of genes known to be frequently hypermethylated in human cancers [8], whereas Arai et al. reported that DNAm alterations in individual patients match in paired adN and tumoral tissues [9]. Moreover, evidence for a gradual increase in epigenetic variability in the sequence of N, adN, and tumoral renal tissues was found [9]. A target-specific comparison of N and adN to differentiate between cell-intrinsic and extrinsically driven DNAm has been carried out by our group for *SFRP1, TBR1, ANKRD34B*, and *ZIC1*, finding both age-dependent and -independent DNAm or, using the terminology described above, both cell-intrinsic and -extrinsic DNAm [7,10,11]. The latter three genes have been identified previously as candidates for age-dependent methylation by biometric analysis of the adN subset of the TCGA KIRC data [12]. Notably, alterations in the DNAm of loci of *ANKRD34B, TBR1*, and *ZIC1* are in line with the results of an epigenome wide association study demonstrating distinct DNAm in adN in breast cancer, as well as the results from Arai et al. describing similarities between the DNAm in adN and tumoral renal tissues [9,13]. DNAm in adN is often designated as a field defect and has been reported for several human adN [14-17]. However, the question of whether these methylation changes demonstrate a rather stochastic characteristic or represent early systematic epigenetic alterations supporting or permitting additional molecular changes with subsequent clonal evolution, such as predicted by the epigenetic stem cell theory of cancer [1], has been sparsely addressed thus far. In a discovery study considering 450K CpG sites, Teschendorff et al. compared the methylation in 50 N and 42 adN breast cancer tissues and found variability in approximately 7,300 CpG sites.

Following biometric reduction, a set of 4,047 candidate loci was evaluated for the epigenetic discrimination of 18 N and 70 adN samples, revealing an area under curve (AUC) of 0.84, providing evidence that non-stochastic methylation occurs in tissues exposed to cancer risk factors. Teschendorff et al. made use of a high-dimensional statistical approach using a number of predictors far exceeding the number of samples to be predicted. In contrast, we asked whether alterations in the 18 CpG sites in *ANKRD34B, NHLH2, TBR1*, and *ZIC1* and the 4 CpG sites in the long interspersed nuclear element (LINE-1) sequence, which is used as a surrogate for detection of genome-wide methylation alterations, are capable of discriminating 342 N and 151 adN renal samples, supporting the hypothesis of the presence of a non-stochastic field defect methylation phenotype in renal adN. Here, we show that renal N and adN can be discriminated with a statistically robust median AUC of 0.81 using DNAm information from 22 CpG sites, indicating the existence of a systematic DNAm field defect in adN from tumor-bearing kidneys.

## 2. Materials and Methods

### 2.1 Identification of candidate loci

This study made use of the measurement of 22 CpG loci annotated to four genes: *ANKRD34B, NHLH2, TBR1*, and *ZIC1*. The LINE-1 repetitive sequence motif was also used as a surrogate marker for global DNAm [18]. The gene-associated candidate loci were identified by biometric analysis of the TCGA KIRC HM450K data and evaluated for age-dependent and age-independent methylation as described previously [10,11,19].

### 2.2 Study design and tissue cohorts

Though candidate loci annotated to the ANKRD34B and ZIC1 genes were part of a previous study [11], the potential association between the DNAm of *NHLH2, TBR1*, and LINE1 loci and the age of tissue donors was measured using a cross-sectional study design with the cohort of N samples. The DNAm of loci in the N and adN cohorts were compared to estimate associated odds ratios (ORs) in a case (adN cohort) - control (N cohort) study. Supervised and unsupervised partitioning was carried out in the united N and adN cohorts. The N cohort consisted of 342 normal autopsy tissues (median age at death 57 years, range 0-98 years), whereas the adN cohort was made up of 151 samples from patients with a median age of 66 years (35-91 years). Of 342 N samples, 119 (35%) and 209 (61%) were derived from female and male donors, respectively. For adN samples, the corresponding numbers were 51 (34%) and 95 (63%) males and females, respectively. Any missing data in the DNAm measurements and covariates age and sex are specified in Table S1.

### 2.3 Analysis of DNA methylation

Histological and molecular evaluation of tissue samples subjected to DNA extraction, bisulfite conversion of DNA, primer design, and subsequent pyrosequencing protocols has been described previously [10,20,21]. Primer sequences for pyrosequencing of candidate loci of *TBR1, NHLH2, ANKRD34B*, and *ZIC1*, as well as the genomic localization of measured CpG sites were specified previously [10,11,19]. LINE1 methylation analysis was carried out according to Park et al. [18].

### 2.4 Statistical analysis

All statistical analyses were carried out in R version 3.6.1 and R-Studio® software using program libraries as specified below [22,23]. Pearson correlation analysis was performed to analyze the CpG site-specific association of DNAm and age. The individual relationship between the independent CpG site-specific DNAm predictor variables, each adjusted by age and sex, and the dichotomous dependent outcome variable of N (control) or adN (case) was exploratively investigated by logistic regression analyses. The selection of an unsupervised classification method for analysis of DNAm in the combined N and adN sample cohorts was carried out using the vcd package [24]. The achievable cluster stability as measured by the Jaccard index was tentatively assessed by the application of various clustering algorithms together with Euclidean or Manhattan distance calculations, the variation of cluster numbers (k=2-10), and each 100-fold bootstrapped repetition of each partitioning. Mosaic plot analysis was used for analysis and presentation of the deviations in the expected proportions of N and adN in k-means partitioning using 2-7 centroids in the clustering [25]. Heat map presentation of the k-means clustering was achieved using the ComplexHeatmap package [26.]

Supervised non-linear classification analysis was carried out using the R tidymodel framework with 1,000 runs for the generation of random splits of tissues into training and test cohorts and random forest classification using default model parameters [27,28]. For each run, the variable importance and receiver operator characteristic (ROC) curve analysis of test cohort classification was performed, and the median AUC, estimated interval of 95% percentiles, and corresponding p-values are presented. Mann-Whitney U statistics were applied to estimate the significance of the AUC being different from the null hypothesis.

## 3. Results

### 3.1 Relationship between DNA methylation and age in normal autopsy tissues

Age-related methylation of CpG loci of *ANKRD34B* and *ZIC1* has been demonstrated previously, but the corresponding analyses of the *LINE1, NHLH2*, and *TBR1* loci were supplemented in this study [11]. Pearson’s correlation analyses showed that 21 out of 22 loci exhibited a significant association between DNAm and age, but inspection of correlation plots and coefficients of correlation revealed heterogeneous results (Figure 1 and Table 1). Therefore, all of the four CpG sites of LINE1, as well as CG1 and CG2 in *NHLH2* clearly show absent or weak relationships between DNAm and age (Figure 1). In contrast, fairly strong positive associations (R> 0.75) were observed for one of the *ANKRD34B* loci and three of the *TBR1* loci, and moderate relationships (R ≈ 0.6) were found for six of the *ANKRD34B* sites, two of the *NHLH2* sites, and each one of the *TBR1* and *ZIC1* CpG sites (Table 1).

**Table 1.** Correlation analysis of the relationship between tissue donor age and relative methylation of CpG sites in normal tissue samples.

| CpG-site | N | R | R<sup>2</sup> | p |
| --- | --- | --- | --- | --- |
| ANKRD34B_CG1 | 326 | 0.755 | 0.571 | 0.000 |
| ANKRD34B_CG2 | 326 | 0.730 | 0.534 | 0.000 |
| ANKRD34B_CG3 | 326 | 0.446 | 0.199 | 0.000 |
| ANKRD34B_CG4 | 326 | 0.714 | 0.509 | 0.000 |
| ANKRD34B_CG5 | 326 | 0.693 | 0.480 | 0.000 |
| ANKRD34B_CG6 | 326 | 0.698 | 0.488 | 0.000 |
| ANKRD34B_CG7 | 326 | 0.617 | 0.380 | 0.000 |
| LINE1_CG1 | 326 | 0.140 | 0.020 | 0.012 |
| LINE1_CG2 | 326 | 0.049 | 0.002 | 0.381 |
| LINE1_CG3 | 326 | 0.367 | 0.135 | 0.000 |
| LINE1_CG4 | 326 | 0.264 | 0.070 | 0.000 |
| NHLH2_CG1 | 319 | 0.385 | 0.148 | 0.000 |
| NHLH2_CG2 | 319 | 0.397 | 0.158 | 0.000 |
| NHLH2_CG3 | 319 | 0.610 | 0.372 | 0.000 |
| NHLH2_CG4 | 319 | 0.648 | 0.420 | 0.000 |
| TBR1_CG1 | 320 | 0.634 | 0.402 | 0.000 |
| TBR1_CG2 | 319 | 0.827 | 0.683 | 0.000 |
| TBR1_CG3 | 319 | 0.865 | 0.747 | 0.000 |
| TBR1_CG4 | 319 | 0.882 | 0.777 | 0.000 |
| ZIC1_CG1 | 306 | 0.391 | 0.153 | 0.000 |
| ZIC1_CG2 | 306 | 0.626 | 0.392 | 0.000 |
| ZIC1_CG3 | 306 | 0.451 | 0.204 | 0.000 |

Results of Pearson's correlation analyses are presented. N, number of normal tissue samples; R, Pearson's coefficient of correlation; p, p-value. Correction for multiple testing was not carried out.

**Figure 1.**
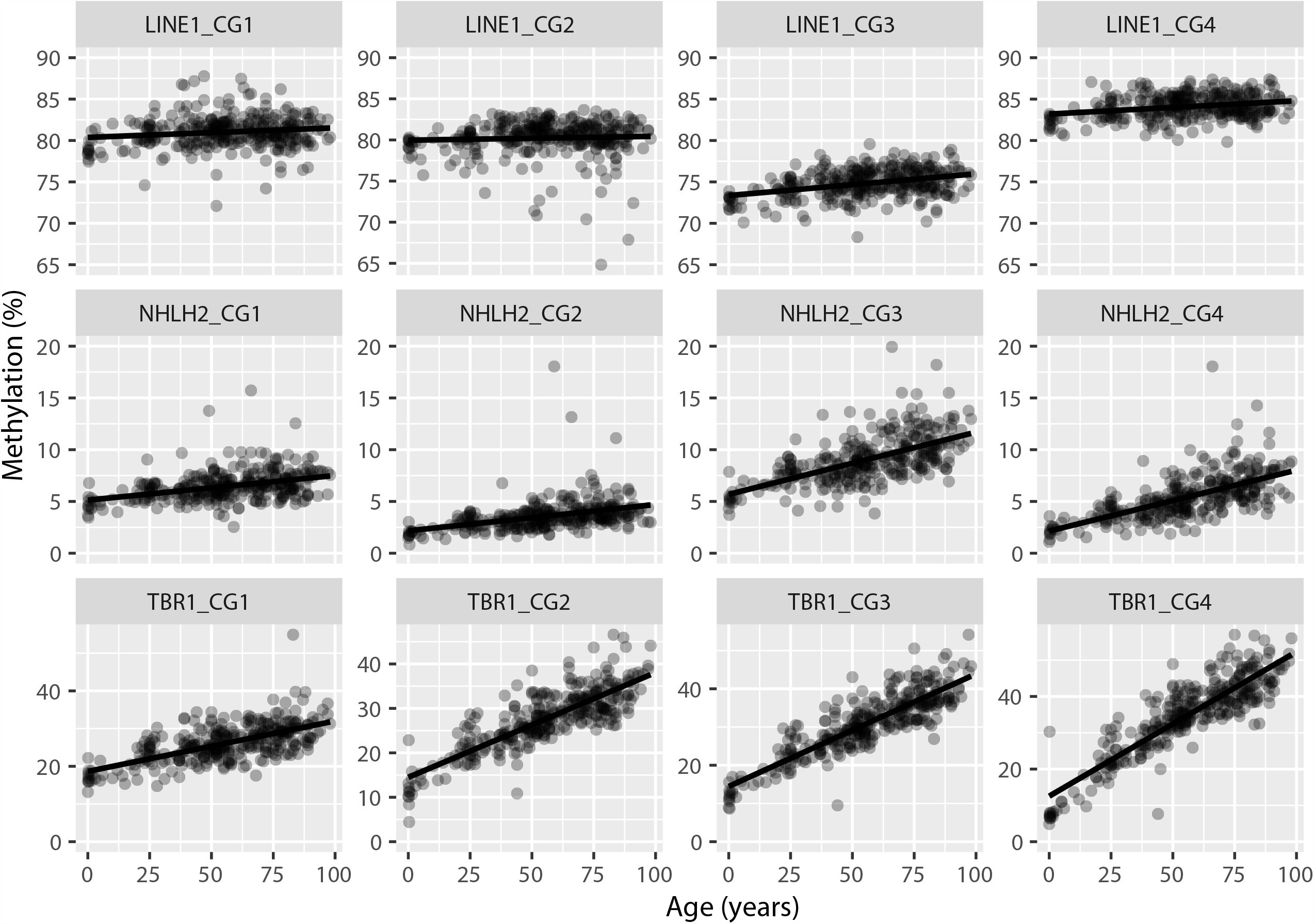
Age-related CpG site-specific methylation of the repetitive LINE1 sequence motif and loci in *NHLH2* and *TBR1* in normal renal tissues. Relative methylation (%) and the linear regression line are shown for normal tissue samples from donors aged 0-98 years. Results for the corresponding correlations analyses are shown in Table 1.

### 3.2 Relationship between DNA methylation and age in normal tumor-adjacent tissues

In previous methylation analyses of the ANKRD34B and ZIC1 genes, we found that DNAm of loci in adN show deviations that are not explainable by the age of the patients, indicating the presence of additional cancer risk comprising factors affecting DNAm, also designated as extrinsic DNAm [2,11]. Analogous supplemental analysis of the loci of *LINE1, NHLH2*, and *TBR1* using visual evaluation of the age-methylation relationship in adN samples and methylation prediction channels calculated from the age-methylation relationship in N samples demonstrated candidate deviations in DNAm in the CG2, CG3, and CG4 loci of LINE1, as a substantial number of samples exhibited relative methylation below the lower boundary of the expected methylation interval (Figure 2, row 1, blue filled squares). In contrast, rare hypermethylation was detected for loci of the *NHLH2* gene, and a more frequent but comparatively low deviation to higher methylation values was found for *TBR1* methylation in adN (Figure 2, rows 2 and 3, red filled squares). Bivariate logistic regression analysis using age and sex as covariates demonstrated significant hypomethylation of CG2-CG4 loci in the LINE1 assay (ORs 0.76-0.80, all p-values < 0.001), a significant but low risk contribution by age, and statistical independence of patient sex (Table 2, Figure 3). Higher ORs of 1.08 and 1.09, potentially indicating hypermethylation of loci in the adN and N comparisons, were detected for CG1 and CG3 of *NHLH2*, but only had borderline significance (p=0.09 and 0.06). The *TBR1* loci methylation comparison in N and adN showed potential hypermethylation of the CG4 locus (OR=1.05, p=0.02), giving no evidence of a significant contribution of the covariates age and sex in the bivariate logistic regression model (Table 2, Figure 3).

**Table 2.**
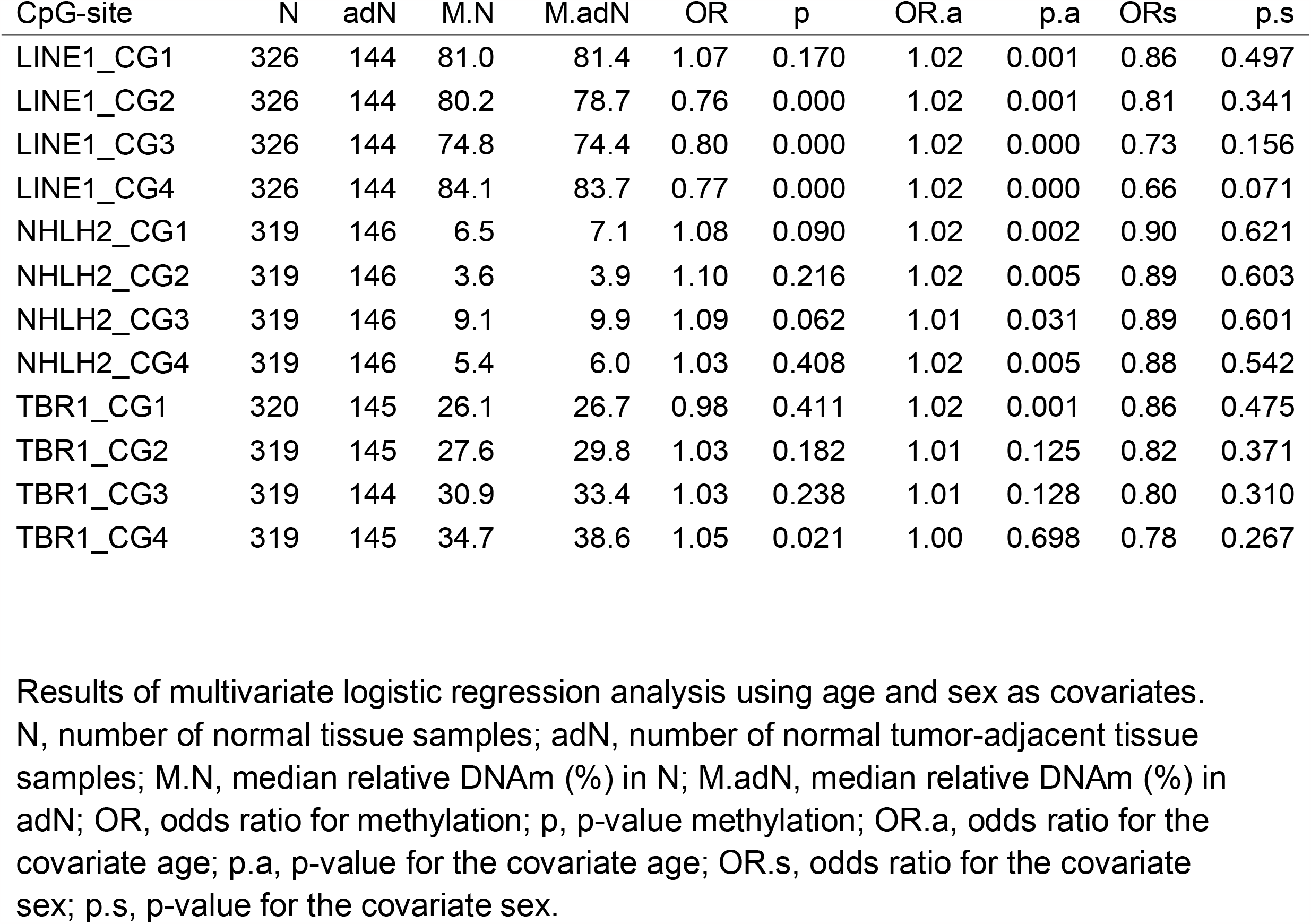
CpG-specific comparison of DNA methylation in normal and tumor-adjacent histopathological normal renal tissues.

| CpG-site | N | adN | M.N | M.adN | OR | p | OR.a | p.a | ORs | p.s |
| --- | --- | --- | --- | --- | --- | --- | --- | --- | --- | --- |
| LINE1_CG1 | 326 | 144 | 81.0 | 81.4 | 1.07 | 0.170 | 1.02 | 0.001 | 0.86 | 0.497 |
| LINE1_CG2 | 326 | 144 | 80.2 | 78.7 | 0.76 | 0.000 | 1.02 | 0.001 | 0.81 | 0.341 |
| LINE1_CG3 | 326 | 144 | 74.8 | 74.4 | 0.80 | 0.000 | 1.02 | 0.000 | 0.73 | 0.156 |
| LINE1_CG4 | 326 | 144 | 84.1 | 83.7 | 0.77 | 0.000 | 1.02 | 0.000 | 0.66 | 0.071 |
| NHLH2_CG1 | 319 | 146 | 6.5 | 7.1 | 1.08 | 0.090 | 1.02 | 0.002 | 0.90 | 0.621 |
| NHLH2_CG2 | 319 | 146 | 3.6 | 3.9 | 1.10 | 0.216 | 1.02 | 0.005 | 0.89 | 0.603 |
| NHLH2_CG3 | 319 | 146 | 9.1 | 9.9 | 1.09 | 0.062 | 1.01 | 0.031 | 0.89 | 0.601 |
| NHLH2_CG4 | 319 | 146 | 5.4 | 6.0 | 1.03 | 0.408 | 1.02 | 0.005 | 0.88 | 0.542 |
| TBR1_CG1 | 320 | 145 | 26.1 | 26.7 | 0.98 | 0.411 | 1.02 | 0.001 | 0.86 | 0.475 |
| TBR1_CG2 | 319 | 145 | 27.6 | 29.8 | 1.03 | 0.182 | 1.01 | 0.125 | 0.82 | 0.371 |
| TBR1_CG3 | 319 | 144 | 30.9 | 33.4 | 1.03 | 0.238 | 1.01 | 0.128 | 0.80 | 0.310 |
| TBR1_CG4 | 319 | 145 | 34.7 | 38.6 | 1.05 | 0.021 | 1.00 | 0.698 | 0.78 | 0.267 |

Results of multivariate logistic regression analysis using age and sex as covariates. N, number of normal tissue samples; adN, number of normal tumor-adjacent tissue samples; M.N, median relative DNAm (%) in N; M.adN, median relative DNAm (%) in adN; OR, odds ratio for methylation; p, p-value methylation; OR.a, odds ratio for the covariate age; p.a, p-value for the covariate age; OR.s, odds ratio for the covariate sex; p.s, p-value for the covariate sex.

**Figure 2.**
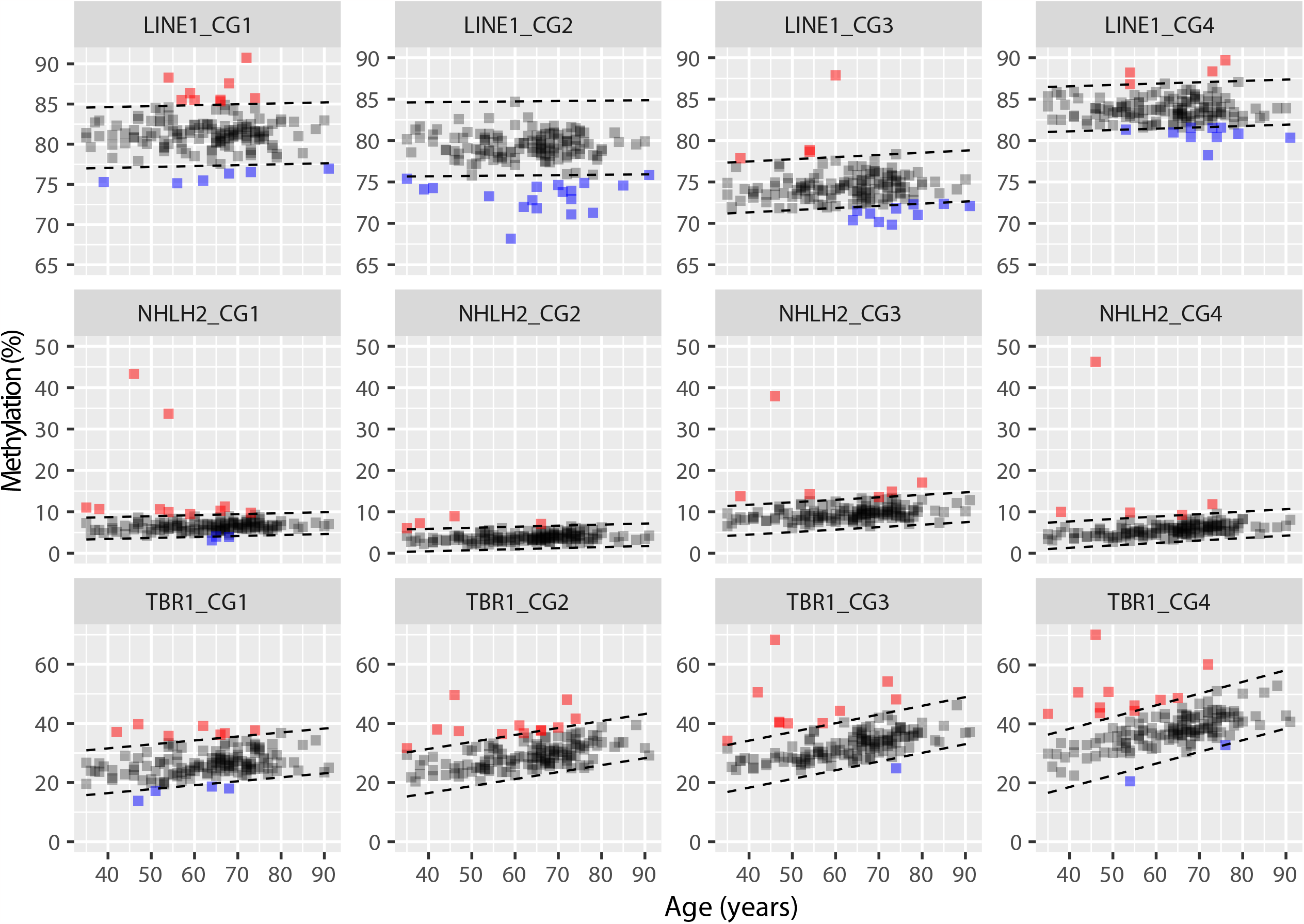
Age-related CpG site-specific methylation of the repetitive LINE1 sequence motif and loci in *NHLH2* and *TBR1* in normal tumor-adjacent renal tissues. The relative methylation (%) of tissues within the 99% methylation prediction interval as defined by normal renal tissues is shown as grey filled squares, whereas relative hypermethylation and hypomethylation are presented as red and blue filled squares.

**Figure 3.**
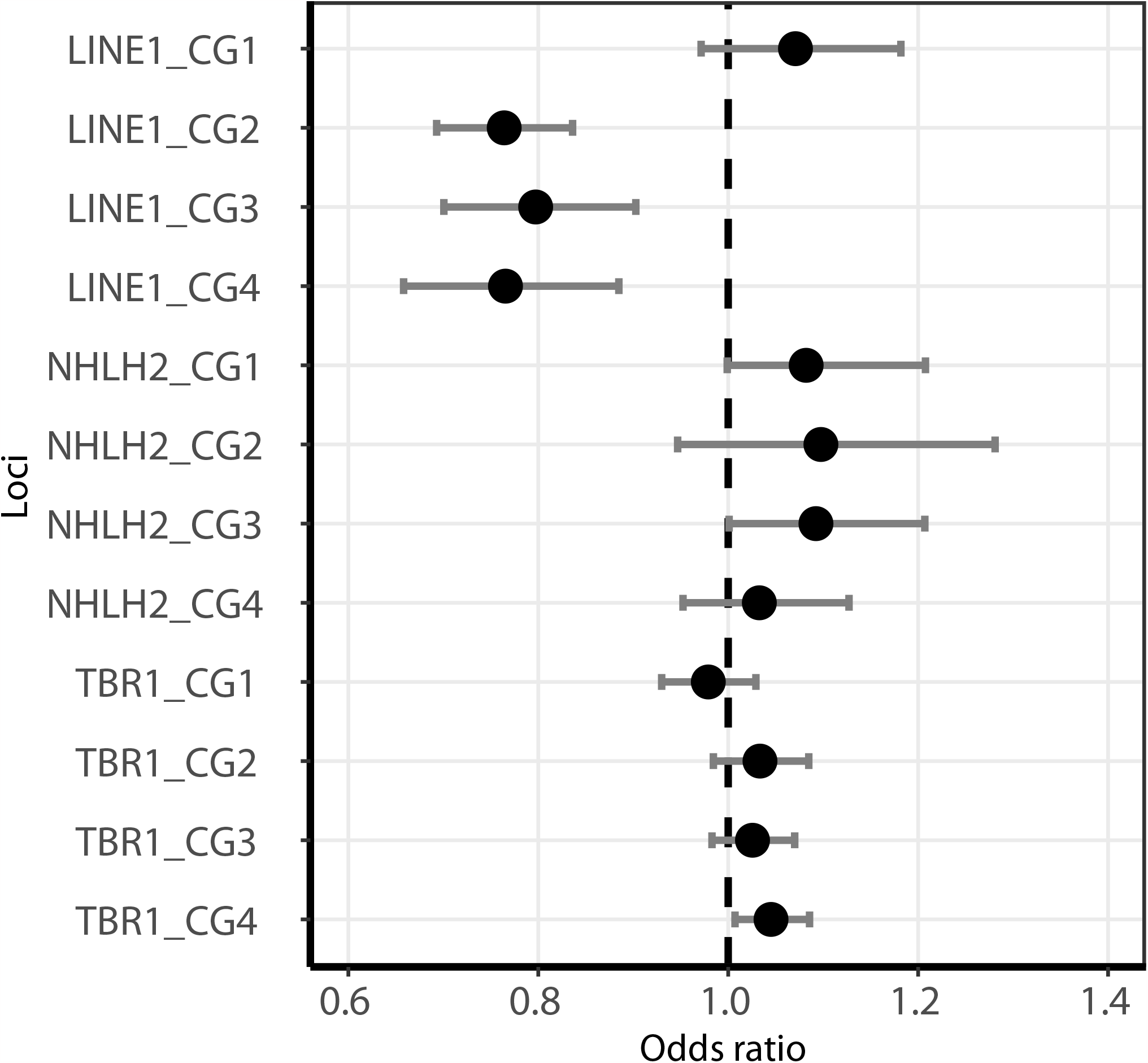
Odds ratios from multivariate logistic regression for the age- and sex-adjusted comparison of CpG site-specific DNAm in normal and normal tumor-adjacent tissues. Also shown are the 95% confidence intervals.

### 3.3 Unsupervised cluster analysis of candidate DNA methylation in normal and tumor-adjacent normal renal tissues

To analyze whether the combined DNAm information from the 22 candidate CpG sites may be informative for discrimination of N and adN, we evaluated various algorithms for unsupervised partitioning using Euclidean or Manhattan distance calculations and assessed the results using the Jaccard index to estimate cluster stability following multiple bootstrapped runs. Highly stable clusters were obtained for k-means partitioning using the Manhattan distance calculation and presets of two, five, or six centroids for clustering (Figure 4a). Jaccard indices of approximately 0.8-0.95 were obtained, indicating high stability of the corresponding clusters in bootstrap runs. Whether stable clustering allows discrimination of tissues was evaluated in Mosaic plot analyses. Significant associations were found for the partitioning into five and six clusters (Figure 4b, c). The separation of tissues into five groups resulted in small clusters 1 and 5, showing an obvious imbalance of the adN and N groups (p=5.4×10^−7^, Figure 4b). The partitioning into six tissue clusters revealed clusters 1, 3, and 5, with significant under- or over-representation of N or adN (p=1.9×10^−7^, Figure 4c). Notably, the most stable clustering of two clusters (Jaccard indices of 0.81 and 0.93) did not show significant discrimination of N and adN (p=1). Visual inspection of the heat map presentation of the partitioning into six clusters showed that either homogeneous clusters were comparatively small and exhibited significant imbalance of N and adN (clusters 1, 3, 5), or were large, such as clusters 2 and 3, and despite showing some degree of homogeneity of methylation patterns, no evidence of a significantly changed proportion of N and adN was provided (Figure 3d).

**Figure 4.**
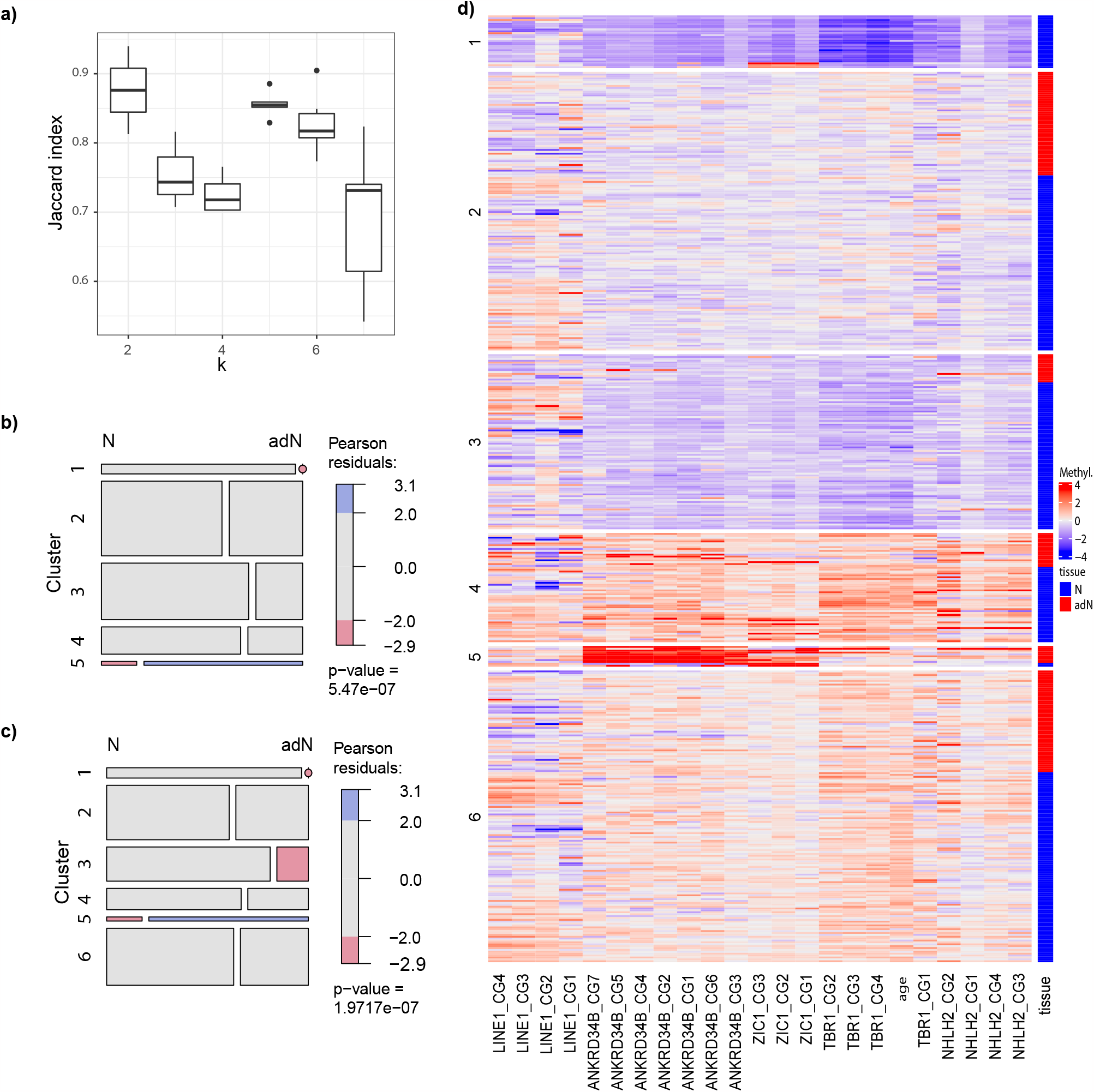
Unsupervised partitioning analysis of DNAm in normal and tumor-adjacent normal tissues. **a)** Box plot presentation of the distribution of Jaccard indices obtained after cluster stability analysis of multiple bootstrapped k-means partitionings using k=2 – 7 cluster centers. **B)** Mosaic plot analysis of the k-means partitioning for five cluster centers as presented in (a). Blue boxes indicate over-represented tissue states in a cluster (Pearson residuals > 2), whereas red boxes correspond to under-represented tissue states (Pearson residuals < −2), providing statistical evidence that clustering is not independent from the tissue state (p = 5.4 × 10^−7^). **c)** Mosaic plot analysis of the k-means partitioning for six cluster centers, providing statistical evidence that this different clustering is not independent from the state of tissues (p = 2 × 10^−7^). **d)** Heatmap presentation of unsupervised partitioning of CpG site-specific DNAm using k-means clustering for six cluster centers. Scaled relative methylation (Methyl.) and normal tissues (N, blue) or normal tumor-adjacent tissues (adN, red) were color-coded as stated in the key.

### 3.4 Supervised statistical classification analyses for the discrimination of normal and normal tumor-adjacent renal tissues

Unsupervised analyses suggested the presence of partially informative methylation data permitting the identification of small tissue-specific clusters, but a large portion of the tissues, though stably clustered, apparently did not allow discrimination of N and adN. Therefore, we analyzed whether the application of the random forest algorithm as a comparatively robust non-linear decision tree method could achieve robust discrimination of both tissues based on the available candidate methylation information. To level out the possible effects of initial random data splitting into training and test subsets, the whole process, including random data splits, random forest model generation, and model evaluation, was repeated 1,000-fold. As a result, we obtained a distribution of ROC curves visually demonstrating a considerable distance of all curves from the diagonal line representing the random classifier, indicating that classification is achieved with good diagnostic efficiency independent from the initial random data splits of training and test subsets (Figure 5). We observed a median AUC for classification of the independent test group of 0.81 (estimated 95% CI 0.74-0.86) and a median p-value 2.7 × 10^−9^ (95% CI 5.5 × 10^−6^ - 8.7 × 10^−13^). The distribution of ROC curves and a representative ROC curve closely matching the median diagnostic parameters is provided in Figure 5a and b. Variable importance analysis for all runs identified CG2 of LINE1, age, CG3 of LINE1, and CG2 of *ANKRD34B* as the overall most important model variables (Figure 5c). Notably, contributions above the overall median importance level were identified in at least one CpG site for all markers except *ZIC1*. To estimate whether the random forest analysis is biased in resulting diagnostic parameters that should be detectable by results showing a systematic deviation from the diagonal line, we carried out 1,000 runs of the complete classification process using random data for each of the independent variables. As a result, we observed a median AUC of 0.50 with an estimated 95% CI of 0.40-0.61 and median p-value of 0.48 (Figure 5d, e). Moreover, analysis of variables of importance showed homogenous values for all metric variables varying around the overall mean importance (Figure 5f).

**Figure 5.**
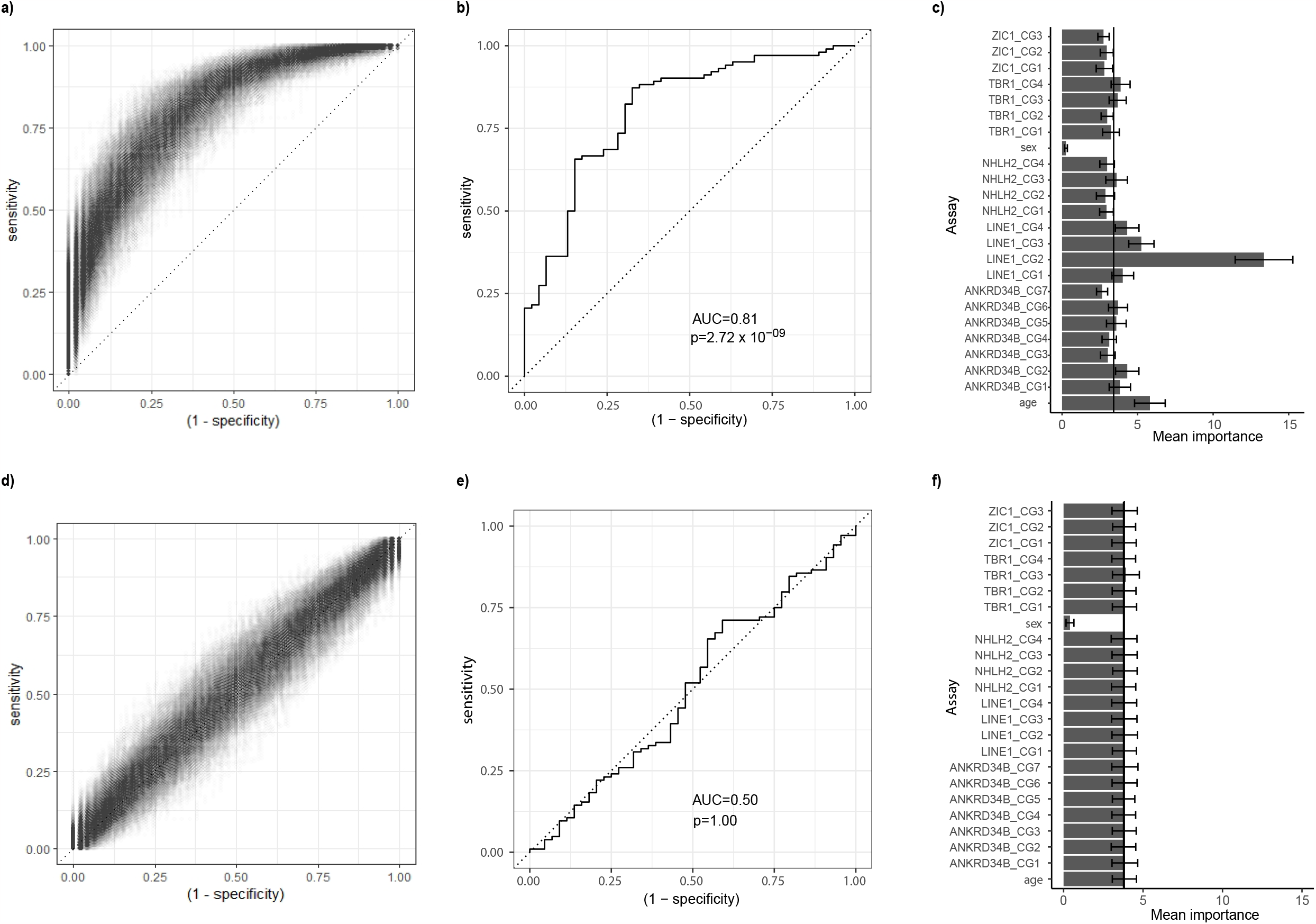
Supervised classification analysis of normal renal tissues using the random forest method. **a)** ROC-AUC analysis of 1,000 random splits of training and test data for random forest classification model generation and evaluation. Only vertices of AUC step curves were used for plotting, applying alpha blending. **b)** AUC analysis of a representative single run closely matching the median diagnostic parameters of multiple runs as shown in (a). **c)** Variable importance analysis of the multiple random forest model generation showing mean importance indices and single standard deviation for the error bars. **d)** ROC-AUC analysis using random data generated for each variable and 1,000 runs as described in (a). **e**) Representative single-run AUC analysis for median AUC values obtained in random data analysis. **f)** Variable importance analysis as described in (c) using random data.

## Discussion

We recently described both age-dependent and age-independent DNAm of loci of *ANKRD34B* and *ZIC1* in normal renal tissues, providing evidence that additional risk factors other than age accelerate DNAm in normal kidney tissue, likely because it is exposed to thus far unknown cancer risk factors. Interestingly, a corresponding analysis of loci of *TBR1* in a subset of our tissue cohort revealed obesity as a candidate factor for accelerated DNAm in adN. Furthermore, considering that loci of *NHLH2* and *TBR1* were previously identified as being associated with metastatic disease progression in RCC and demonstrated increased epigenetic variability in our normal tissue cohorts ([10,19] and unpublished data), the question was raised as to whether these DNAm alterations combined with data on the LINE1 marker as a surrogate for global methylation are sufficient to allow differentiation of N and adN samples.

Though analysis for age-dependent DNAm of *ANKRD34B* and *ZIC1* loci in N already demonstrated an age-dependent or cell-intrinsic increase in our previous study, the corresponding investigation of the LINE1, *NHLH2*, and *TBR1* sequences demonstrated heterogeneous results. Two of the four sites in *NHLH2* and all *TBR1* loci exhibited age-related methylation when using a coefficient of correlation of R > 0.6 as the criterion. Considering that we recently found that *NHLH2* loci exhibit both tumor- and metastatic tissue-specific hypermethylation that participates in a renal metastasis-associated methylation signature, both the finding of additional age-dependent DNAm of *NHLH2* loci and the corresponding results for *TBR1* loci point to cell-intrinsic epigenetic alterations in normal renal tissues likely persisting in tumoral tissues and possibly affecting the clinical course of RCC [29]. In contrast, only one locus of LINE-1 had a coefficient of correlation of R>0.3, indicating no relevant association with age overall. Data for LINE-1 global methylation in solid human tissues, including kidney, are relatively sparse. Though a consensus exists that LINE-1 normally exhibits tumor-specific hypomethylation in human solid cancers, such was not found for RCC in an early analysis [30,31]. However, alteration of LINE-1 methylation in noncancerous esophageal mucosae and white blood cells has been associated with the cancer risk factors age and smoking [32,33].

Accelerated DNAm independent of age has been detected previously for the *ANKRD34B* and *ZIC1* loci, but our complementary analysis of the LINE1, *NHLH2*, and *TBR1* loci revealed a heterogeneous methylation characteristic [11]. In the case of *NHLH2*, corresponding loci visually demonstrated increased evidence of epigenetic variability independent from age, appearing mostly as hypermethylation, with a comparably low frequency and only reaching borderline significance in bivariate logistic regression analysis. In contrast, DNAm of LINE1 sites presented with a considerable number of mostly hypomethylated outliers when visually analyzing the age-DNAm relationship in adN. The corresponding age-adjusted statistical comparison of N and adN for age-independent risk contribution identified comparable high odds for hypomethylation (1.32 per 1% change in DNAm) in the LINE1-CG2 (OR=0.76 for hypermethylation). Whether the comparatively high age-independent variability in LINE-1 methylation observed for the CG1-CG3 sites in adN can be attributed to specific extrinsic life style factors, such as smoking, is a relevant question that remains to be clarified. Overall, significant alteration of DNAm not explainable by age was observed in at least one locus of each of the LINE1, *ANKRD34B, ZIC1*, and *TBR1* methylation assays, but the ORs and borderline significance indicate that two loci of *NHLH2* could also represent age-independent cancer risk loci provided a larger cohort were available for analysis. Therefore, our analysis adds three LINE1 loci and potentially two of the *NHLH2* loci to the limited list of current candidate sites showing DNAm associated with extrinsic cancer risk factors in normal kidney tissue. We questioned whether the combined methylation information on all 22 of the available CpG sites would allow discrimination of N and adN, which would support the view of a “stand alone” DNAm characteristic of renal adN and defining an epigenetic field effect differentiating cancer risk-exposed normal tissue from normal counterparts.

Our unsupervised classification analysis of DNAm data using k-means clustering provided highly stable clusters for the 2, 5, or 6 cluster center presets, but significant enrichment of N or adN was only obtained in smaller clusters for the separation of five and six clusters, respectively. Mosaic plot analysis and visual inspection of the corresponding heatmap analysis revealed that clusters 1 and 3 had mostly low methylation values for the *ANKRD34B, NHLH2, TBR1*, and *ZIC1* sequences to be associated with significant overrepresentation of N. In contrast, only the small cluster 5 exhibited high DNAm values for the *ANKRD34B* and *ZIC1* loci and consisted predominantly of adN. On the other hand, large clusters 2 and 6, characterized by a median to moderately higher methylation of all CpG sites beside LINE-1 loci, demonstrated no significant imbalance of tissue types. We hypothesize that the limited tissue discrimination in cluster analysis could be a consequence of only subsets of tissues with increased epigenetic variability of our candidate loci being identified. Consequently, systematic identification of loci showing age-independent increases in methylation variability in appropriate N vs. adN comparisons would be expected to improve the partitioning of tissues. However, assumptions and presets made for unsupervised partitioning may interfere with the efficient discrimination of tissues. Therefore, we questioned whether random forest classification, as a non-linear robust supervised partitioning method, is capable of more efficient discrimination of tissues.

Following 1,000-fold randomization of the initial data split into training and test subsets, we obtained a distribution of ROC curves clearly differing in all of the analysis runs from the bisector line, indicating diagnostic indifference, with an AUC of 0.5. Thus, random forest analysis showed that N and adN can be discriminated in a robust manner with a good median diagnostic efficiency and, moreover, that this distinction can be made solely on the basis of patient age and the relative degree of methylation of the candidate loci, defining adN of the kidney as a distinct tissue from normal healthy renal counterparts. This finding is in line with the concept of an epigenetic field defect identifying tissues with increased cancer risk. Importance analysis of the random forest analysis demonstrated an above median relevance of three of the LINE-1 sites, age, and at least one of each the other gene-annotated CpG sites, providing evidence that all sequences contribute to decision-making in the algorithm. The likely prominent role of the CG2 site of LINE-1 in variable importance analysis may be reflected by an unusually high epigenetic variability in the analyses of age-dependency of adN, indicating increased informativity of this site.

Notably, though making use of only 23 predictor variables, the results of our classification analyses (summarized by an AUC of 0.81) compare to the diagnostic efficiency of the previous comparison of breast N and adN that reported an AUC of 0.84 and the use of 4,047 variables. Also taking into account our previous findings, according to which the methylation of both *NHLH2* and *TBR1* loci is associated with metastatic progression of tumors, we assume that the distinct methylation observed in the renal adN samples likely does not show a stochastic behavior. In contrast to the breast tissue study, our approach did not make use of high-dimensional methylation array data subject to possible statistical problems in view of the comparatively small test and training cohorts [34]. A strength of our study is that the number of samples to be predicted is substantially higher than the number of predictors (n/p ratio > 20), exceeding the ratio of approximately 0.02 used in the breast tissue study. Correspondingly, no evidence of any bias or overfitting of the classification process was found as a result of our runs using randomized data (median AUC=0.50). On the other hand, our study likely could benefit from additional markers exhibiting an age-independent increase in variability in the adN group, underlining the need for systematic candidate marker discovery following a genome-wide comparison of renal adN and N. Another limitation of this study is the lack of tissue donor-specific risk data, which can possibly be mitigated as measurable surrogates increasingly appear.

The superior importance of LINE-1 loci in the random forest importance analysis could hypothetically be due to the observation that LINE-1 hypomethylation in normal esophageal mucosae is associated with smoking, which is also an important cancer risk factor for RCC [32]. Therefore, the important role of LINE-1 methylation in our classification of renal adN samples clearly requires further analysis.

Notably, our results may also be relevant for the development of tissue-based epigenetic clocks. Though most epigenetic candidate clocks have been developed by using DNA extracted from blood or its components, they apparently suffer from consistent results for the detection or prediction of several solid cancers, possibly due to alterations in DNAm showing substantial tissue specificity and limiting the efficiency of surrogate tissue measures, such as of blood cells [2,35]. On the other hand, our current random forest model is capable of detecting adN, which is equivalent to the indirect detection of RCC or precancerous tissues. Yet, any application of our methylation signature clearly suffers from the lack of available solid renal tissues, currently ruling out a non-invasive assay. However, this problem could be potentially solved in the medium term, as recent analyses demonstrated that, in principle, all types of normal renal cells can be found in urine and, therefore, an at least hypothetical test set-up for an epigenetic tissue-based clock for RCC prediction is conceivable [36].

## Conclusions

In conclusion, histopathologically normal tissues adjacent to renal tumors exhibit a DNAm field defect, permitting distinction from healthy normal tissue by a robust, low-dimensional, statistical classification approach. This may represent the basis for a future solid normal tissue-based epigenetic clock predicting RCC risk and potential progression of the disease.

