## Supplementary material for "Epigenetic field defect discriminates normal tissues from healthy and tumor-bearing kidneys": Suppl Table 1

**Table S1.** Assay and tissue-specific numbers for the available data on the DNA methylation (meth), age, and sex of tissue donors

| Assay | Tissue | N.meth | N.age | N.sex | NA.meth | NA.age | NA.sex |
| --- | --- | --- | --- | --- | --- | --- | --- |
| ANKRD34B | N | 342 | 326 | 328 | 0 | 16 | 14 |
| ANKRD34B | adN | 151 | 146 | 146 | 0 | 5 | 5 |
| LINE1 | N | 342 | 326 | 328 | 0 | 16 | 14 |
| LINE1 | adN | 149 | 146 | 146 | 2 | 5 | 5 |
| NHLH2 | N | 335 | 326 | 328 | 7 | 16 | 14 |
| NHLH2 | adN | 151 | 146 | 146 | 0 | 5 | 5 |
| TBR1 | N | 336 | 326 | 328 | 6 | 16 | 14 |
| TBR1 | adN | 150 | 146 | 146 | 1 | 5 | 5 |
| ZIC1 | N | 322 | 326 | 328 | 20 | 16 | 14 |
| ZIC1 | adN | 137 | 146 | 146 | 14 | 5 | 5 |

Values are the number of measurements. N, normal tissue samples; adN normal tumor-adjacent tissue samples; NA, not available.
